# Identifying dynamic, partially occupied residues using anomalous scattering

**DOI:** 10.1101/642686

**Authors:** Serena Rocchio, Ramona Duman, Kamel El Omari, Vitaliy Mykhaylyk, Zhen Yan, Armin Wagner, James C. A. Bardwell, Scott Horowitz

## Abstract

X-ray crystallography is generally used to take single snapshots of a protein’s conformation. The important but difficult task of characterizing structural ensembles in crystals is typically limited to small conformational changes, such as multiple side-chain conformations. A crystallographic method was recently introduced that utilizes Residual Anomalous and Electron Density (READ) to characterize structural ensembles encompassing large-scale structural changes. Key to this method is an ability to accurately measure anomalous signals and distinguish them from noise or other anomalous scatterers. This report presents an optimized data collection and analysis strategy for partially occupied iodine anomalous signals. Using the long wavelength-optimized beamline I23 at Diamond Light Source, the ability to accurately distinguish the positions of anomalous scatterers with as low as ~12% occupancy is demonstrated. The number and position of these anomalous scatterers are consistent with previous biophysical, kinetic and structural data that suggest the protein Im7 binds to the chaperone Spy in multiple partially occupied conformations. This study shows that a long-wavelength beamline results in easily validated anomalous signals that are strong enough to be used to detect and characterize highly dynamic sections of crystal structures.

**Synopsis:** Structural studies on partially occupied, dynamic protein systems by crystallography are difficult. We present methods here for detecting these states in crystals.

## 1. Introduction

In crystallography, anomalous scattering is commonly used to help solve the phase problem (Hendrickson, 2014). A second, less well-utilized aspect of anomalous scattering is its ability to selectively label and identify residues of interest. Crystallographers can use anomalous maps to pinpoint metal ions (Handing *et al.*, 2018) or to aid in model building or electron density interpretation (Pflug *et al.*, 2012; Wang *et al.*, 2012). We recently introduced a method called READ that uses anomalous maps to allow the reconstruction of highly heterogeneous conformational ensembles in regions of the protein that are not well ordered enough for traditional model building (Salmon *et al.*, 2018; Horowitz *et al.*, 2016). This method uses selective anomalous labeling with iodo-phenylalanine (pI-Phe) to generate multiple partially occupied iodine anomalous signals in the crystal corresponding to different protein conformations. Ensemble selection techniques (Venditti *et al.*, 2016; Salmon *et al.*, 2018; Horowitz *et al.*, 2016) are then used to create ensembles that are consistent with both the anomalous data and weak electron density data for the dynamic segment(s) of the crystal.

A critical step in the READ process is the correct identification of weak anomalous signals (Horowitz, Salmon, *et al.*, 2018; Wang, 2018). Selective labeling requires tunable X-ray sources to achieve high levels of anomalous scattering for particular atoms (Phillips *et al.*, 1976). However, many important anomalous scatterers do not have their K and L absorption edges within wavelength ranges of standard synchrotron beamlines for macromolecular crystallography, resulting in weak anomalous signals. Recently, a novel beamline, I23 at Diamond Light Source, was introduced to collect anomalous data at substantially longer wavelengths than was previously possible. The I23 beamline operates in an in-vacuum sample environment and features a number of other innovations that make it optimal for anomalous data collection (Wagner *et al.*, 2016). Here, we present an optimized data collection strategy for identification of iodine anomalous scattering positions that greatly improves our ability to detect weak and partially occupied states in crystals. Co-crystals of the chaperone Spy with its dynamic client Im7 were used to collect anomalous data for three iodine-containing Im7 mutants. Using multiple wavelengths and collections from different orientations obtained with a multi-axis goniometer, we demonstrate that it is possible to detect the iodine signal even at low occupancy and clearly distinguish it from other anomalous scatterers. The anomalous signals indicate that Im7 binds to Spy in multiple conformations along its concave surface, consistent with findings from other biophysical investigations. The data demonstrate the feasibility of using anomalous scatterers as an approach to understand dynamic systems in crystallography.

## 2. Methods

### 2.1. Crystallization

The super Spy chaperone mutant, Spy H96L (Quan *et al.*, 2014), was used here for co-crystallization experiments, as we found it formed crystals more readily than wild-type Spy protein (Horowitz *et al.*, 2016). Spy-Im7 co-crystals were obtained using the vapor diffusion technique and the crystallization conditions 22–34% PEG 3000, 70–270 mM zinc acetate, and 0.1 M imidazole, pH 8.0. Crystallization was performed using 15–50 mg/ml Spy H96L, with Im7 peptides that represent the 6–26 portion of the Im7 peptide in a 1:1–1:2 ratio. The Im7 peptides (^6^SISDYTEAEFVQLLKEIEKEN^26^) used here for the co-crystallization experiments have a single iodo-phenylalanine substitution in one of the positions 18, 19, and 20, respectively. The Im7 6–26 peptides bind to Spy H96L with a 7.8 ± 0.6 μ*M* dissociation constant, which is similar to the dissociation constant of a partially folded variant of Im7 for wild-type Spy (Fig. S1).

#### 2.1.1. Crystal harvesting

Crystals were cryoprotected by increasing the PEG 3000 concentration up to 35% and flash frozen in liquid nitrogen. Crystals were harvested using LithoLoop sample mounts (Molecular Dimensions, Newmarket, UK) matched to the size of the crystal, glued to copper pins as part of the dedicated I23 sample holder assembly, optimized for cryogenic sample transfer and storage in the beamline vacuum environment.

#### 2.1.2. Data collection

Data were collected at beamline I23 at Diamond Light Source at X-ray energies of 5.2 and 4.5 keV, above and below the iodine L absorption edges (*E*_L(I)_ = 5.188 keV, *E*_L(II)_ = 4.852 keV, *E*_L(III)_ = 4.557 keV) using the semi-cylindrical PILATUS 12M area detector. For each data set, 360° of data were collected with an exposure time of 0.1 sec per 0.1° rotation in inverse beam setting of 20° wedges. Taking advantage of the multi-axis goniometer, data were collected at 5.2 keV at different *κ* and *φ* goniometer angles (Table 1). To identify the remaining anomalously scattering ligands, the zinc and chloride ions, two additional sets of data were collected for the Spy:Im7-L19pI-Phe complex at 2.87 and 2.75 keV.

**Table 1.**
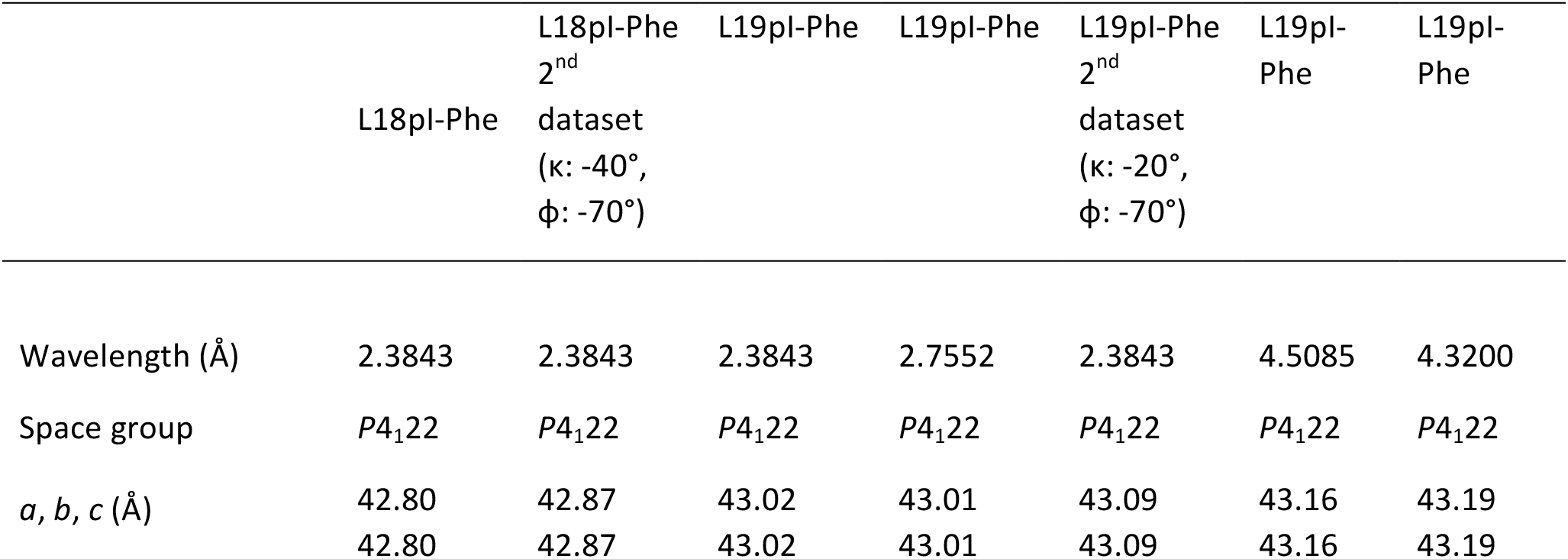
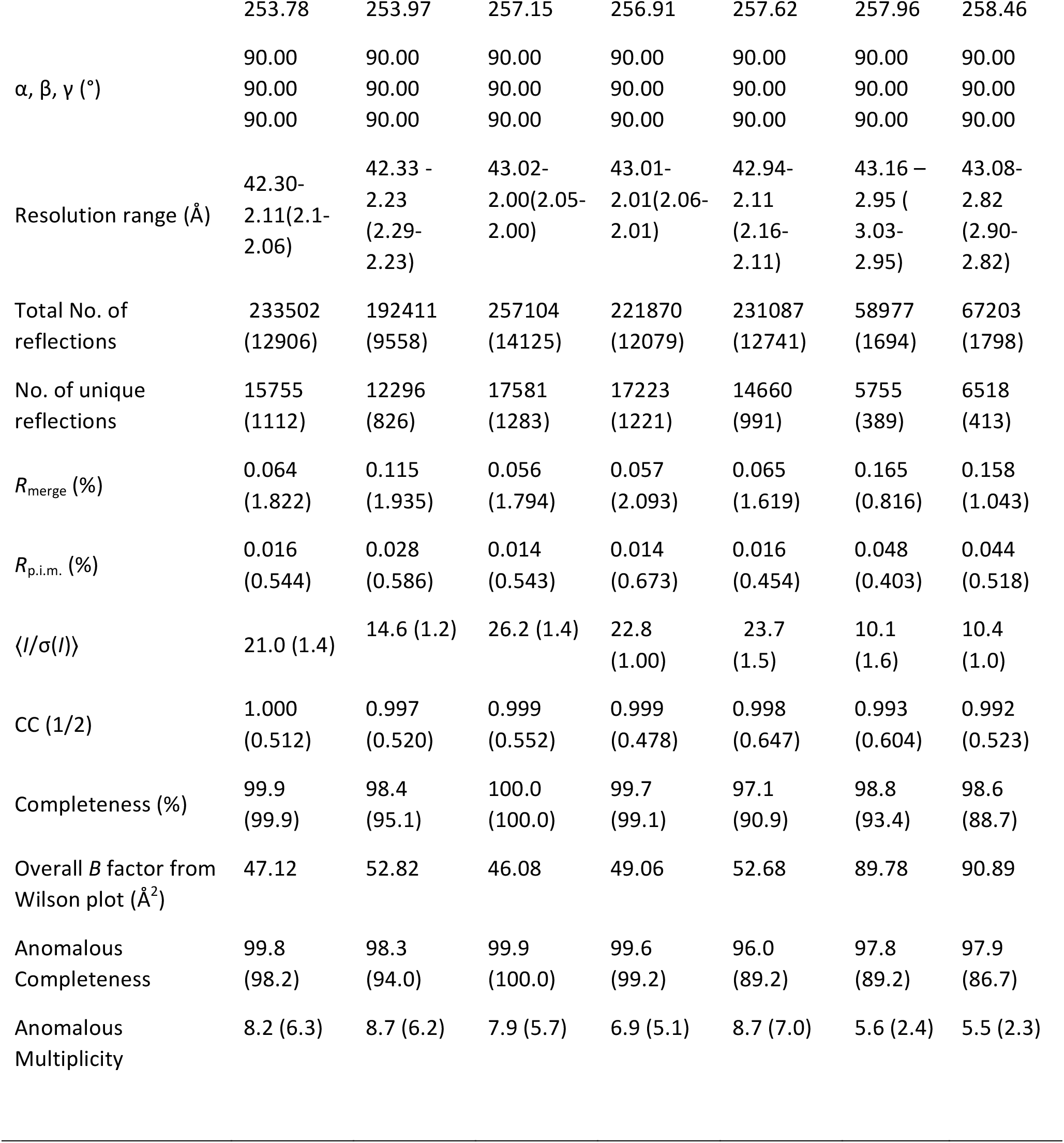
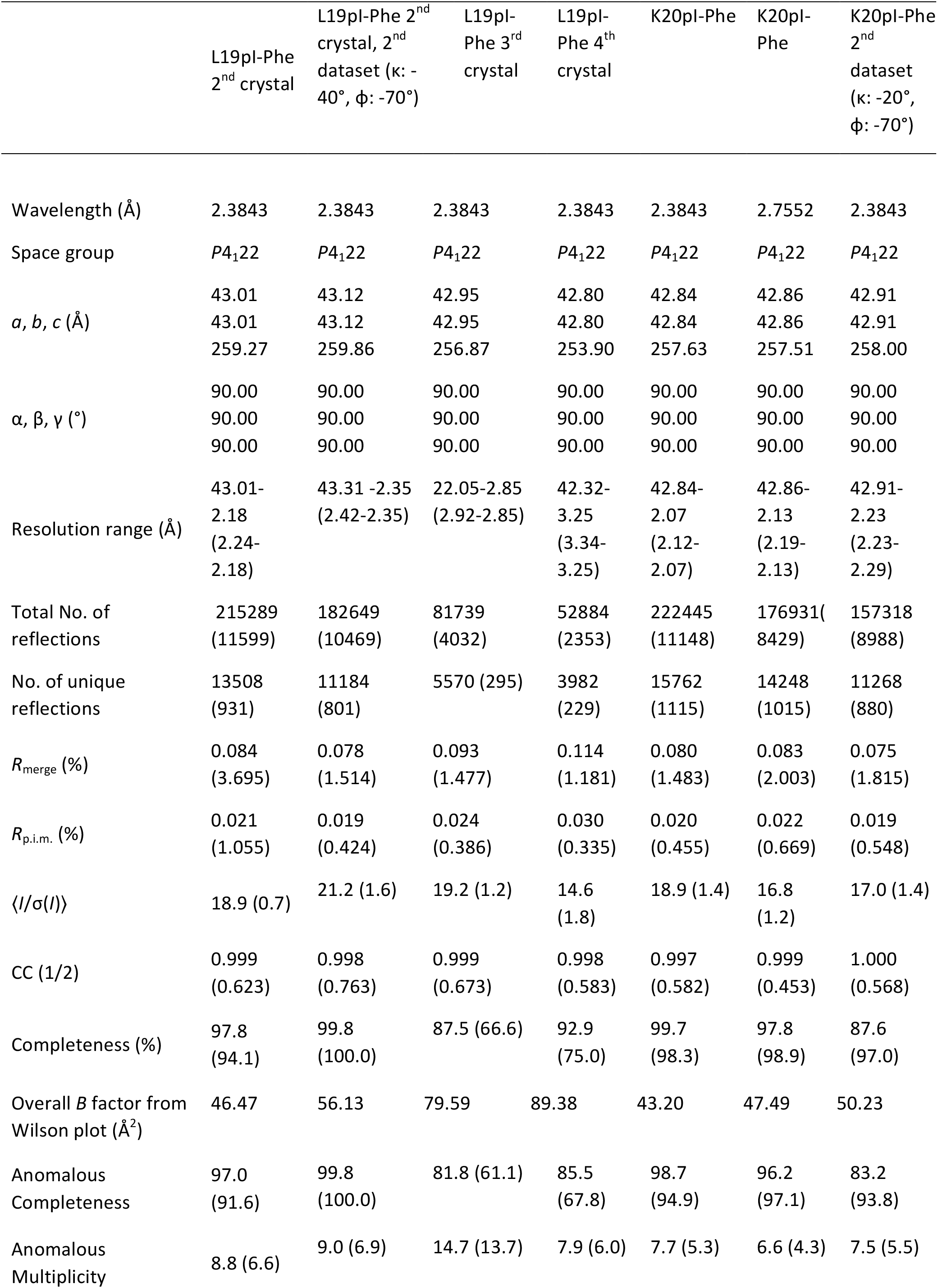
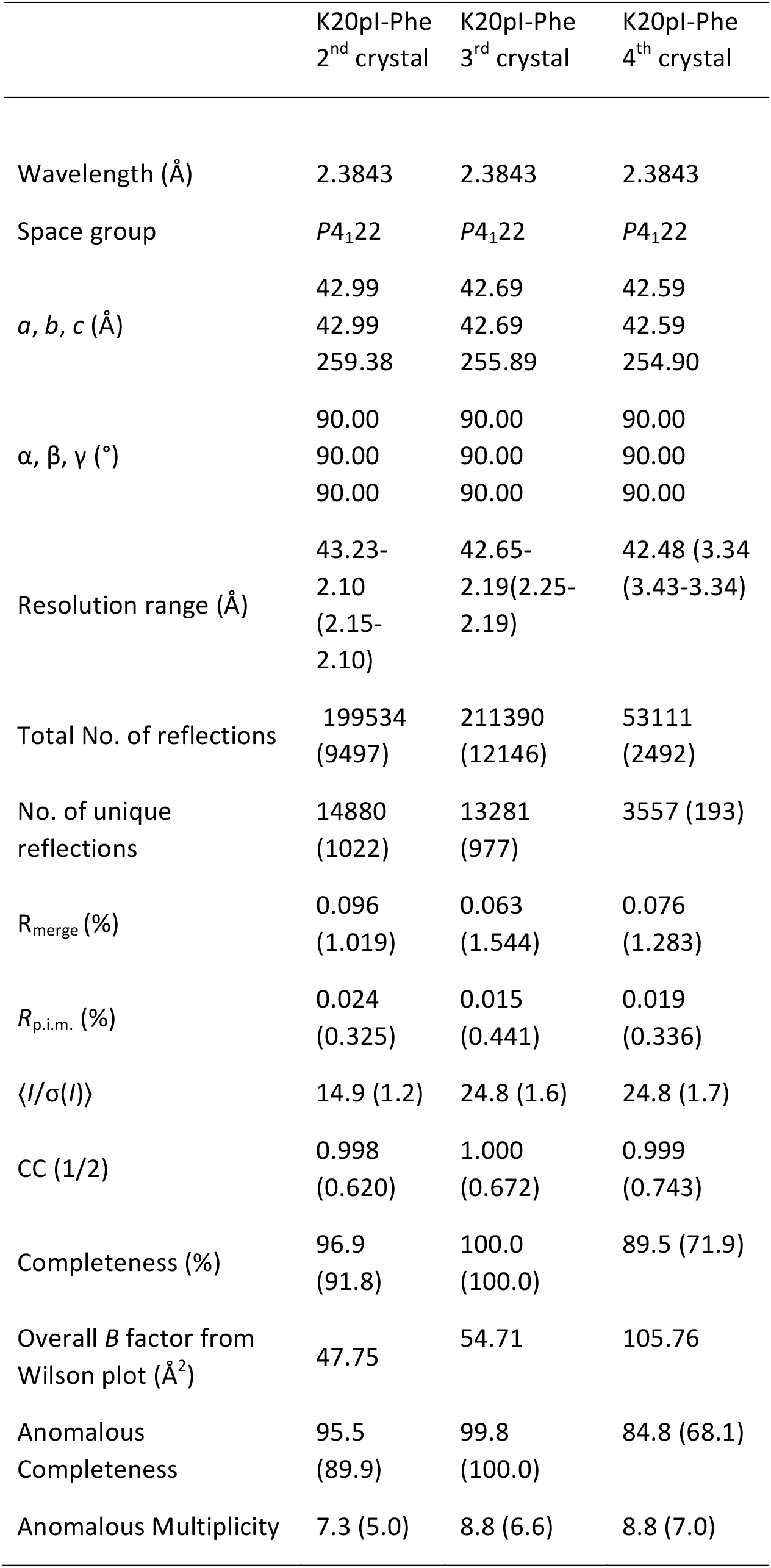
Data Collection and processing. Values for the outer shell are given in parentheses.

**Table 2.** Refinement.

|  | L18pl-Phe | L19pl-Phe | K20pl-Phe |
| --- | --- | --- | --- |
| PDB Code | 6OWX | 6OWZ | 6OWY |
| Resolution range (Å) | 42.79 - 2.06 (2.134 - 2.06) | 42.43 - 2.05 (2.123 - 2.05) | 42.26 - 2.07 (2.144 - 2.07) |
| Completeness (%) | 99.89 (99.87) | 99.94 (100.00) | 99.65 (99.70) |
| CC 1/2 | 1 (0.651) | 0.999 (0.861) | 0.999 (0.691) |
| CC* | 1 (0.888) | 1 (0.962) | 1 (0.904) |
| No. of reflections, working set | 15669 (1516) | 16247 (1538) | 15674 (1511) |
| No. of reflections, test set | 996 (93) | 976 (92) | 939 (91) |
| Final $R_{\text{cryst}}$ | 0.2256 (0.3250) | 0.2150 (0.2778) | 0.2268 (0.3604) |
| Final $R_{\text{free}}$ | 0.2803 (0.3851) | 0.2549 (0.2945) | 0.2724 (0.4299) |
| No. of non-H atoms |  |  |  |
| Protein | 1373 | 1382 | 1403 |
| Ion | 15 | 13 | 13 |
| Ligand | 26 | 19 | 16 |
| Water | 42 | 62 | 58 |
| Total | 1456 | 1476 | 1500 |
| R.m.s. deviations |  |  |  |
| Bonds (Å) | 0.011 | 0.015 | 0.013 |
| Angles (°) | 1.21 | 1.20 | 1.23 |
| Average $B$ factors (Å<sup>2</sup>) | 72.77 | 60.61 | 58.32 |
| Protein | 72.58 | 60.72 | 58.74 |
| Ion | 78.20 | 64.87 | 66.28 |
| Ligand | 88.18 | 63.03 | 55.78 |
| Water | 73.17 | 54.87 | 48.21 |
| Ramachandran plot |  |  |  |
| Most favoured (%) | 95.91 | 95.68 | 94.71 |
| Allowed (%) | 3.51 | 3.70 | 4.71 |

#### 2.1.3. Model building and refinement

Data integration and scaling were performed with *XDS* (Kabsch, 2010) and *AIMLESS* (Evans & Murshudov, 2013), respectively. Anomalous difference Fourier maps were calculated using *ANODE* (Thorn & Sheldrick, 2011). Molecular replacement was performed using the known structure of the Spy-Im7 complex (PDB entry 5wnw) as the search model in *Phenix* (Adams *et al.*, 2010). Model building and refinement were accomplished using *Coot* (Emsley *et al.*, 2010) and *Phenix* (Afonine *et al.*, 2012), respectively. For lowly occupied iodine atoms, harmonic restraints were applied in *Phenix* refinement to prevent them from moving from the anomalous density. Iodine occupancies were estimated by performing refinements with anomalous groups enabled. For occupancy refinement, *B* factors and occupancies were initiated from differing starting positions, with the final occupancies examined to determine whether convergence was achieved, following the protocol described in (Langan *et al.*, 2018). The protein structures were very similar in each case, with the maximum C*_α_* root-mean-square deviation (r.m.s.d.) = 0.41 Å between protein structures.

## 3. Results

### 3.1. Data collection strategy

Spy is one of a group of chaperones that allows clients to fold while remaining continuously bound to the chaperone’s surface (Horowitz, Koldewey, *et al.*, 2018). We have previously observed that the chaperone substrate Im7 exists in a heterogeneous, largely disordered crystallographic ensemble while bound to the chaperone Spy (Horowitz *et al.*, 2016). To refine strategies for detecting anomalous scattering for observing partially occupied dynamic states, we co-crystallized the tight-binding Spy H96L variant with short peptides derived from its client, Im7. These peptides each had a single iodo-phenylalanine substitution at positions 18, 19, and 20, respectively.

Our data collection strategy was geared to determine the positions of these iodines with maximum sensitivity while ensuring that the signals were due to iodine and not from other weak anomalous scatterers present in the crystal. To maximize signal intensity, we selected data collection wavelengths that should give complementary data based on the anomalous scattering peaks and the edges of iodine (Fig. S2). We collected data primarily at *λ* = 2.3843 Å (*E* = 5.2 keV), just above the L(I) absorption edge of iodine, as well as below the L(III) edge at *λ* = 2.7552 Å (*E* = 4.5 keV). At these two energies, the anomalous contributions *f*” to the scattering factor differ by a factor of 3.9 (*f*”_4.5keV_ = 3.42 e- vs. *f*”_5.2keV_ = 13.41 e-). By collecting in this fashion, we reasoned that we should be able to specifically distinguish iodine anomalous signals from other with elements, by analysing the resulting anomalous difference fourier maps. Peaks that are present in the higher energy dataset (above the iodine edge), but absent in the lower energy dataset (below the iodine edge) (*λ* = 2.7552 Å, 4.5 keV) originate from iodine atoms. We further reasoned that one of the best ways to distinguish noise from real signals is to show that the same anomalous signal exists in independent datasets collected at different crystal orientations. We thus collected datasets at *λ* = 2.3843 Å (*E* = 5.2 keV) from different goniometer *κ* angles (*κ* = −40°, *φ* = −70° and *κ* = −20°, *φ* = −70°) so we could compare and stringently determine the number and positions of the iodine peaks. Finally, when possible, we compared data from multiple crystals with the same components to see the reproducibility of the anomalous signals. To help identify zinc and chlorine sites in the crystal, we also collected datasets at *λ* = 4.3200 Å (*E* = 2.87 keV) and *λ* = 4.5085 Å (*E* = 2.75 keV), just above and below the chlorine K absorption edge (*E* = 2.822 keV), respectively. Despite the very long wavelengths, the data quality was good enough to observe the peaks expected for chlorine and zinc ions (Table 3, Fig. S3). To our knowledge, these are the first protein crystallography datasets collected around the chlorine absorption edge.

**Table 3:** Anomalous signals detected for zinc and chlorine anomalous signal peak heights (in σ), from the 2.87 and 2.75 keV ANODE maps collected on the best L19pI-Phe Spy:Im7 crystal.

| *Atom* | **2.87 keV Peak Height** | **2.75 keV Peak Height** |
| --- | --- | --- |
| *Zn 1* | 3.6 | 5.6 |
| *Zn 2* | 4.6 | 5.0 |
| *Zn 3* | 5.0 | 6.1 |
| *Zn 4* | 4.2 | 4.7 |
| *Zn 5* | 5.3 | 7.8 |
| *Zn 6* | 6.9 | 9.6 |
| *Zn 7* | 14.0 | 10.8 |
| *Cl 1* | 12.0 | 0.3 |
| *Cl 2* | 11.0 | 1.1 |

### 3.2. L19pI-Phe demonstrates multiple binding poses of Im7 on Spy

From the three peptides tested, anomalous maps of L19pI-Phe show the highest number of partially occupied anomalous signals that we can attribute to iodine. In addition to the anomalous density attributed to zinc, chlorine, and methionine, there are four distinct anomalous signals above 6 *σ* in the *ANODE* anomalous maps at 5.2 keV (Fig. 1*a* and Table 4). Of these four peaks, three (iodine number 1, 2, and 4) are very well validated by additional maps, as discussed below, whereas the remaining peak is less well supported (Fig. 1*c*). When refined from multiple different starting occupancies and *B* factors, these iodines displayed both low occupancy and high temperature factors, with occupancies estimated to range from 0.12 to 0.29, and *B* factors ranging from 89 to 133 Å^2^ (Table 4).

**Table 4:** Iodine anomalous signal peak heights (in σ), occupancies and B-factors (Å^2^) from the 5.2 keV ANODE maps for the best crystal of each Spy:Im7 complex.

| *Spy:Im7* | **Peak Height** | **Occupancy** | **B-Factor** |
| --- | --- | --- | --- |
| *L18pl-Phe* | 26.6 | 0.27 | 50.6 |
| *L19pl-Phe 1* | 8.9 | 0.22 | 89.3 |
| *L19pl-Phe 2* | 9.8 | 0.29 | 132.7 |
| *L19pl-Phe 3* | 6.3 | 0.23 | 115.8 |
| *L19pl-Phe 4* | 6.4 | 0.12 | 106.4 |
| *K20pl-Phe* | 32.4 | 0.48 | 65.2 |

**Figure 1:**
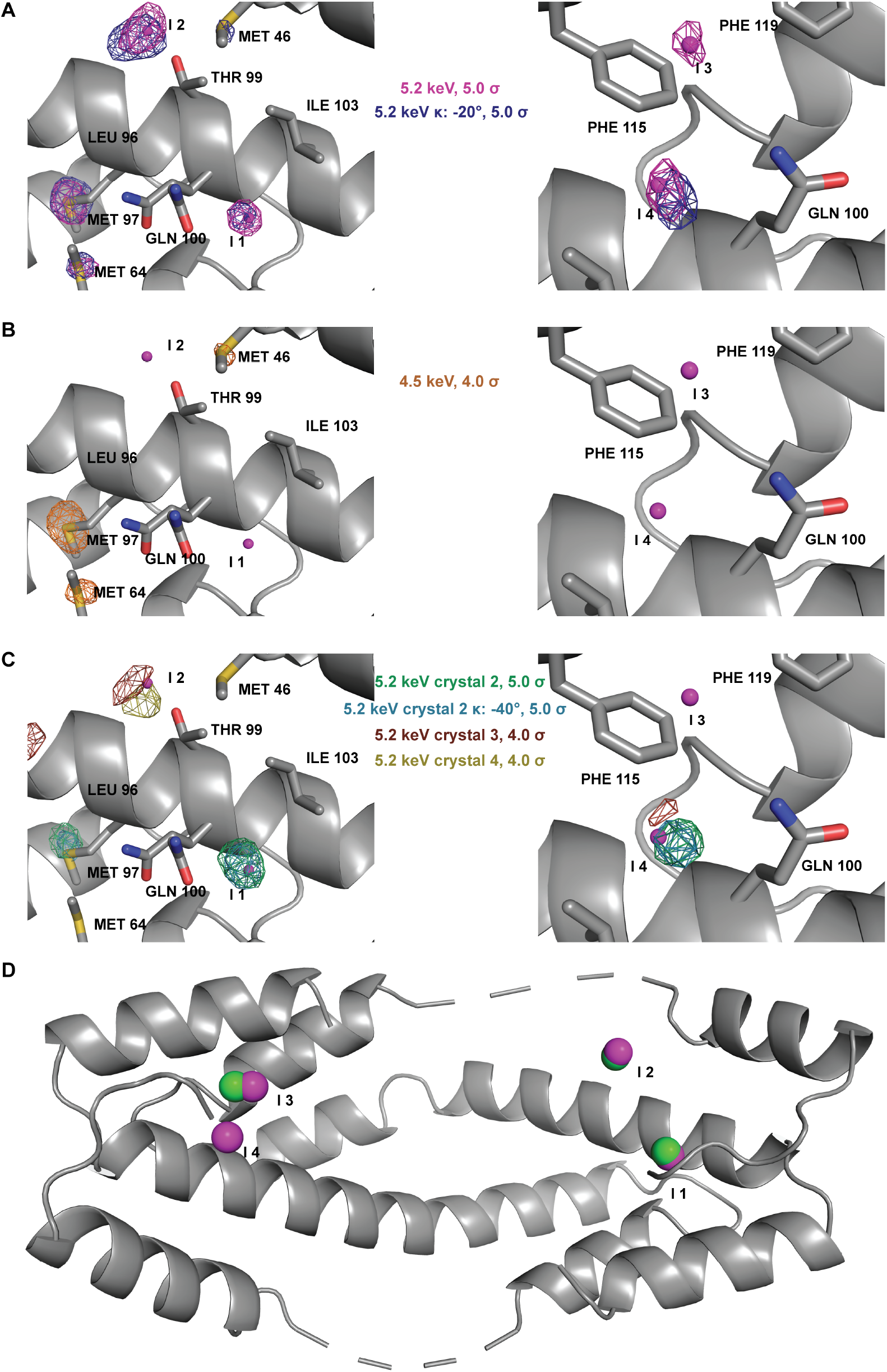
Anomalous maps and positions of the four iodine signals in L19pI-Phe crystals, displayed as purple spheres in each panel, which were defined through the displayed anomalous fourier maps. (A) ANODE anomalous difference fourier maps from 5.2 keV data collected at two different goniometer kappa angles, contoured at 5.0 σ. (B) ANODE anomalous map from 4.5 keV contoured at 4 σ. (C) ANODE anomalous maps from data collected from replicate crystals, contoured at 5.0 σ (crystal 2) and 4 σ (crystal 3 and 4). (D) Overlap of L19pI-Phe iodine positions (magenta) detected here with closest iodine signals from previous data (green) (Horowitz *et al.*, 2016).

To check if these signals are attributable to iodine, we collected an additional dataset on the same crystal at 4.5 keV, below the iodine L edges. At this energy, we would expect that the anomalous signal derived from iodine would dramatically decrease, whereas the change in anomalous signal from methionine, zinc, and chlorine would be negligible. The *ANODE* anomalous maps from this energy confirm that the non-iodine scatterers continue to display strong anomalous signals, whereas the signals from the four putative iodines disappear (Fig. 1*b*). This experiment indicates that the four anomalous signals are from the Im7 peptides and not from other anomalous scatterers. Moreover, the high *σ* values of the anomalous signals, the smallest of which was detected at 6.3 *σ*, make it unlikely that these signals represent noise. As a counterexample, the highest peaks not attributable to known anomalous scatterers, which hence could be noise, have *σ* values of 4.2 in the same map. However, a rigorous way to exclude the possibility that the anomalous signals do represent noise would be to determine if they are present in the same positions in an additional dataset collected at the same energy (at 5.2 keV) but using a different goniometer *κ* angle (*κ* = −20°). Anomalous signals that appear in the same positions are highly likely to be true anomalous signals. It is worth mentioning that after a complete set of data, the intensity of the anomalous signal will decrease due to radiation damage. Therefore, only the signals with the highest intensity are detected after multiple additional data collections on the same crystal. The *ANODE* anomalous maps from this additional data collection show that three of the anomalous positions from the first data collection are preserved in the data collected at the different *κ* angle (Fig. 1*a*).

Finally, to test whether these anomalous positions are reproducible in other L19pI-Phe crystals, we collected additional 5.2 keV datasets from three other L19pI-Phe crystals. These crystals diffracted to resolutions of 2.2, 2.9, and 3.2 Å, respectively. In the anomalous map of the crystal that diffracted to 2.2 Å, we were able to identify three different iodine signals that overlap with signals detected in the highest resolution crystal (Fig. 1*c*). These signals were also present in other data collections (*κ* = −40°) on the 2.2 Å diffracting crystal (Fig. 1*c*). In the crystals that diffracted to 2.9 and 3.2 Å, we only detected one iodine signal in each, but they overlapped well with the strongest anomalous signal from the highest resolution crystal. It is therefore clear that the ability to detect low intensity signals is highly dependent on the crystal and the resolution.

To further examine the anomalous peaks detected here, we also compared these iodine positions to those identified in our previous Spy-Im7 crystallography study in which we had used the same L19pI-Phe peptides with H96L Spy. We found that three of the four iodine anomalous positions detected here were also detected and used in our previous analysis (Fig. 1*d*).

The lowest L19pI-Phe anomalous peak (6.3 *σ* at 5.2 keV) is not present in the anomalous dataset from the crystals diffracting to lower resolution. However, this signal is very close (1.3 Å) to an L19pI-Phe anomalous peak observed in our previous Spy-Im7 crystallography study (Fig. 1*d*). So, although this peak was not well supported by additional datasets within this study, it was detected in our previous study.

### 3.3. Other Im7 peptides suggest additional Im7 binding sites in the crook of Spy’s cradle

The Im7 L18pI-Phe and K20pI-Phe peptides crystallized in complex with Spy show a very different pattern compared to the multiple signals observed for L19pI-Phe. The 5.2 keV *ANODE* anomalous difference maps of 18pI-Phe and 20pI-Phe each show only one distinct anomalous peak. In both cases, the peak heights in the *ANODE* anomalous maps were very strong; the iodine signals were detected at 26.6 and 32.4 *σ* for L18pI-Phe and K20pI-Phe, respectively (Figs. 2 and 3, and Table 4). Both peaks are on the interior of Spy’s cradle, near the flexible loop region.

**Figure 2:**
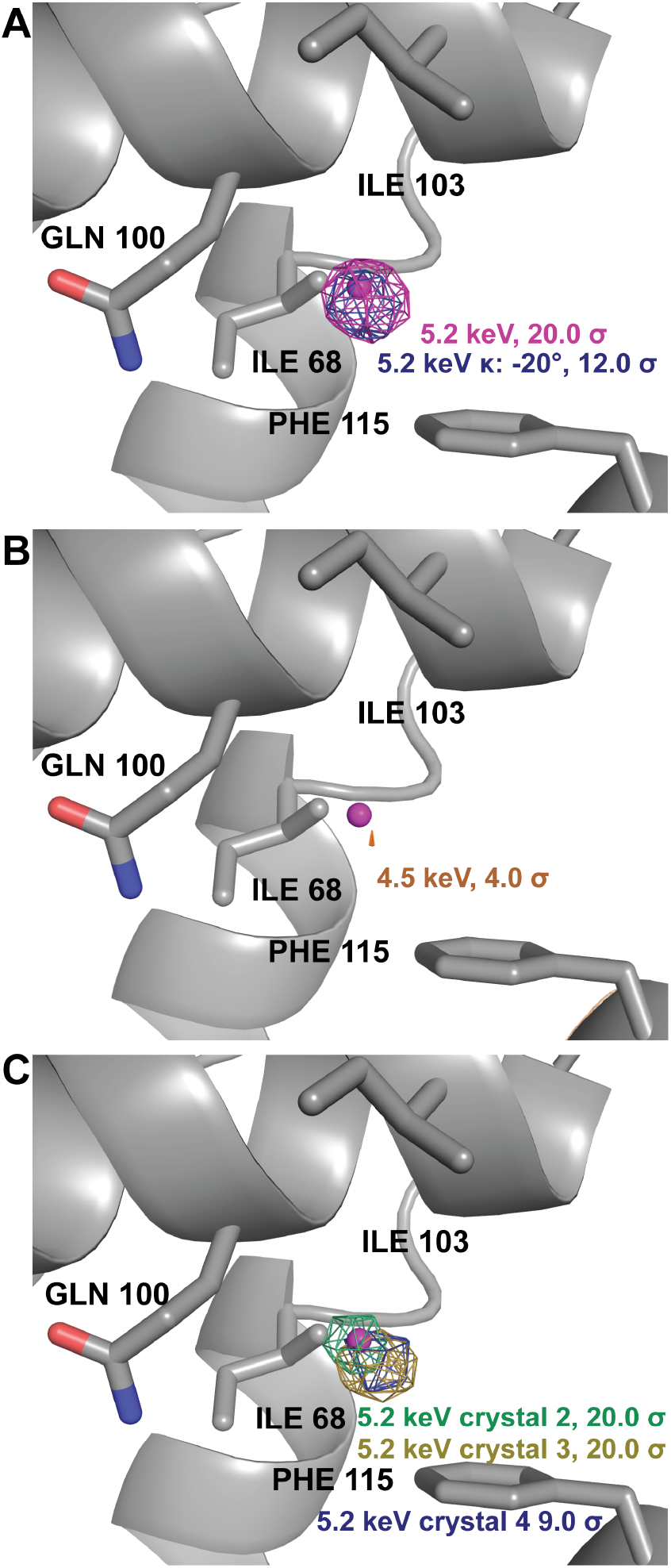
K20pI-Phe anomalous maps, with iodine atom depicted as a magenta sphere. (A) *ANODE* anomalous maps from 5.2 keV datasets collected at two goniometer kappa angles, contoured at 20 σ and 12 *σ*, respectively. (B) *ANODE* anomalous map from data collected at 4.5 keV, contoured at 4.0 *σ*. (C) *ANODE* anomalous maps from data collected 5.2 keV from three additional crystals, contoured at 20 *σ* (crystal 2 and 3) and 9 *σ* (crystal 4).

**Figure 3:**
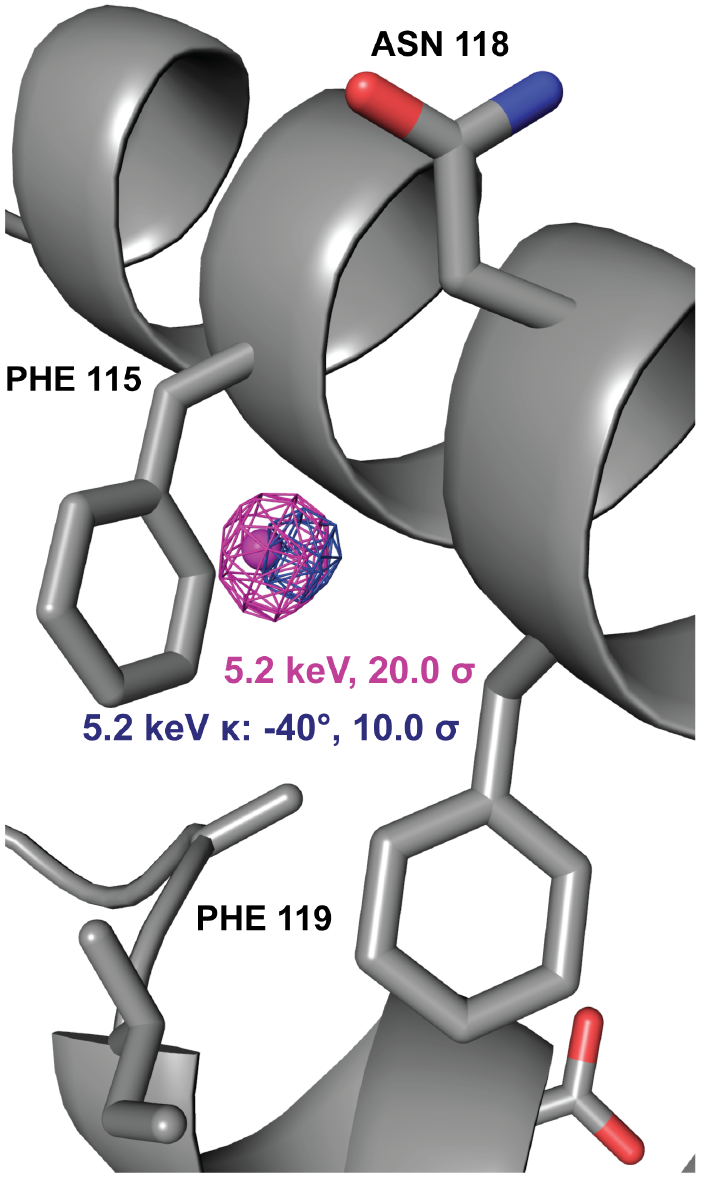
L18pI-Phe *ANODE* anomalous maps collected at 5.2 keV at two different goniometer *κ* angles, with iodine depicted as a magenta sphere.

For K20pI-Phe, a second dataset below the iodine edge at 4.5 keV was collected. The large decrease of the anomalous signal in this dataset confirmed that the position ascribed to iodine was indeed iodine (Fig. 2*b*). To further validate this observation, we collected a third dataset at an additional goniometer *κ* angle (*κ* = −20°) above the iodine edge. This dataset again showed strong anomalous density (16.1 *σ* in the *ANODE* map) at the same position (Fig. 2*a*)), with the peak height reduced by radiation damage. Finally, we collected datasets at 5.2 keV from three additional crystals that showed strong anomalous signals in the same position (26.7, 26.0, and 6.5 *σ* in *ANODE* maps), demonstrating good reproducibility for this signal (Fig. 2*c*). Consistent with the L19pI-Phe data collections, the strength of the iodine anomalous signal detected is highly dependent on the crystal quality. This combination of measurements confirmed that K20pI-Phe produced detectable anomalous scattering from a single position in the crystal, providing the ideal case for determining the position of a partially occupied anomalous scatterer.

Unfortunately, for L18pI-Phe, the single crystal that had diffraction adequate for anomalous analysis degraded during the second 5.2 keV dataset collections. (Fig. 3 and Table 4). This dataset, collected at *κ* = −40°, confirmed the presence of the anomalous signal (Fig. 3), but the 4.5 keV dataset could not be analyzed further due to radiation damage. However, other indicators strongly suggest that this anomalous peak also arises from iodine: (1) The height of the anomalous peak is the second largest observed in this study, and at 26.6 *σ* in the *ANODE* map, is substantially larger than the signals observed from zinc, chlorine, and methionine sulfur, the largest of which is 9.3 *σ*; (2) The position of this anomalous signal is structurally inconsistent with expected zinc or chlorine binding residues, which would be required to enable high enough occupancy binding to produce a strong anomalous signal. For example, this signal does not occur in the vicinity of zinc or chloride binding residues. So although the 4.5 keV dataset was not analyzed for this crystal, the strength of the signal and its chemical environment are inconsistent with it being derived from zinc or chloride, making it likely that the anomalous signal is due to the iodine in the Im7 peptide.

Further support for the L18pI-Phe signal coming from iodine derives from the observation that carbon-iodine bonds are labile and cleavage is expected to occur due to radiation damage (von Schenck *et al.*, 2003). Thus, for anomalous signals of peptide-bound iodine, we would expect to see a decrease in signal intensity as a function of increasing X-ray dose, as the iodine is cleaved from the peptide backbone. This trend is demonstrated not just for L18pI-Phe, but also for K20pI-Phe in Table 5.

**Table 5:** Radiation damage specifically affects iodine anomalous signals. As data collection proceeds, the anomalous signals (in σ from ANODE maps) for both L18pI-Phe and L20pI-Phe drop while that of a nearby methionine sulfur stays constant.

| *Spy:lm7* | **L18pl-Phe** |  | **K20pl-Phe** |  |
| --- | --- | --- | --- | --- |
| *Residue* | Met93 | Iodine | Met93 | Iodine |
| *1<sup>st</sup> 5.2 keV dataset* | 4.5 | 26.4 | 4.9 | 32.5 |
| *2<sup>nd</sup> 5.2 keV dataset* | 4.0 | 10.6 | 4.2 | 16.1 |

The larger peak heights of L18pI-Phe and K20pI-Phe, as compared to each of the L19pI-Phe peaks, suggest a higher level of occupancy at these positions. Anomalous site refinements show that the occupancies of L18pI-Phe and K20pI-Phe are approximately 25% and 48% with *B* factors of approximately 50 and 65, respectively (Table 4). Similarly, the area adjacent to the anomalous signals in L18pI-Phe and K20pI-Phe shows electron density in 2*F*_o_ — *F*_c_ and *F*_o_ — *F*_c_ maps consistent with the presence of a partially occupied, dynamic peptide (Fig. 4). Attempts to perform traditional model building using the residual L18pI-Phe and K20pI-Phe peptide density were unsuccessful due to the weak nature of the electron density, as well as its somewhat amorphous shape, likely due to multiple conformations within the peptide. This density, however, still shows greater residual peptide occupancy than the areas surrounding the multiple L19pI-Phe iodine anomalous peaks, where we failed to see density above background, consistent with the higher occupancy of the single L18pI-Phe and K20pI-Phe positions.

**Figure 4:**
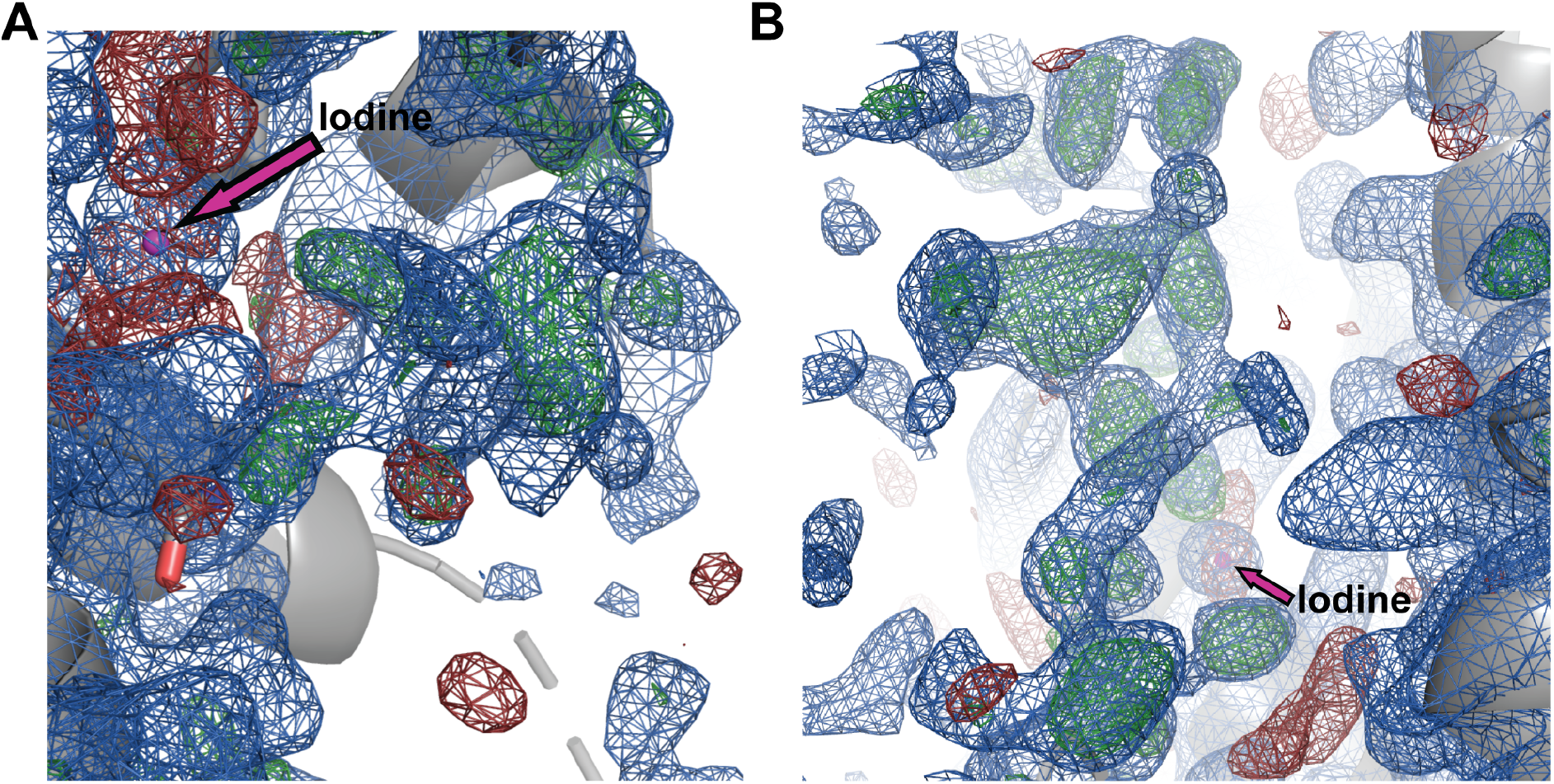
Residual electron density from disordered peptides. (*a*) K20pI-Phe peptide density. (*b*) L18pI-Phe peptide density. 2*F*_o_ — *F*_c_ map is displayed in blue, contoured at 0.6 *σ*, and *F*_o_ — *F*_c_ map is displayed in green and red, contoured at 2.5 *σ*.

## 4. Discussion

Our new, optimized data collection strategy provides an improved method to obtain high quality anomalous signals with less noise contamination; it also verifies the presence of several of the iodine anomalous signals identified in our previous datasets. Combined, the experiments demonstrate that the Im7 peptide binds to Spy in multiple different binding poses, which are detectable using anomalous scattering. Refining the occupancies of the iodine anions suggests that we are able to detect iodine anomalous signals as low as ~12% occupancy, even with a high temperature factor of 106 Å^2^. These lowly occupied states were confirmed via reproducibility through independent data collections at multiple angles and using separate crystals.

Spy-Im7 binding has recently received attention as a model system for understanding chaperone-client interactions (He *et al.*, 2016; He & Hiller, 2018; Salmon *et al.*, 2016). NMR spectroscopy, molecular dynamics simulations, chemical kinetics, and X-ray crystallography have all concluded that client binding to Spy is dynamic and that Im7 can bind to Spy in an array of conformations and poses (He *et al.*, 2016; He & Hiller, 2018; Salmon *et al.*, 2016; Stull *et al.*, 2016; Horowitz *et al.*, 2016). These studies showed that Im7 binding occurs at various sites on the concave surface of the Spy cradle. The work here confirms these findings and provides an avenue to identify binding sites directly at higher sensitivity than was previously possible. Combining the six iodine positions observed from three different iodine-containing peptides, the data reported here confirms that Im7 binds to Spy in multiple binding poses.

The anomalous signals detected from the three peptides suggest an interesting binding pattern (Fig. 5). The peaks for both L18pI-Phe and K20pI-Phe are both very close to separate L19pI-Phe anomalous peaks. This proximity suggests a degree of promiscuity in the binding sites in which Im7 can shift small distances to accommodate pI-Phe binding. Moreover, based on the decreasing peak intensities of the iodine anomalous signals through the data collection (Table 4), we can rule out that these positions are iodine ions that have been cleaved from pI-Phe, as this would have produced the opposite trend. Instead, it is likely that these positions are acceptable binding locations for the iodine that are accessible within the Im7 ensemble in the crystal. These positions are not equivalent, as no iodine density in L18pI-Phe is visible at the K20pI-Phe site, or vice-versa. These measurements confirm the observations that Im7 binds to Spy promiscuously, but with specific anchor points between Spy and Im7 (He *et al.*, 2016; He & Hiller, 2018; Salmon *et al.*, 2016; Stull *et al.*, 2016; Horowitz *et al.*, 2016).

**Figure 5:**
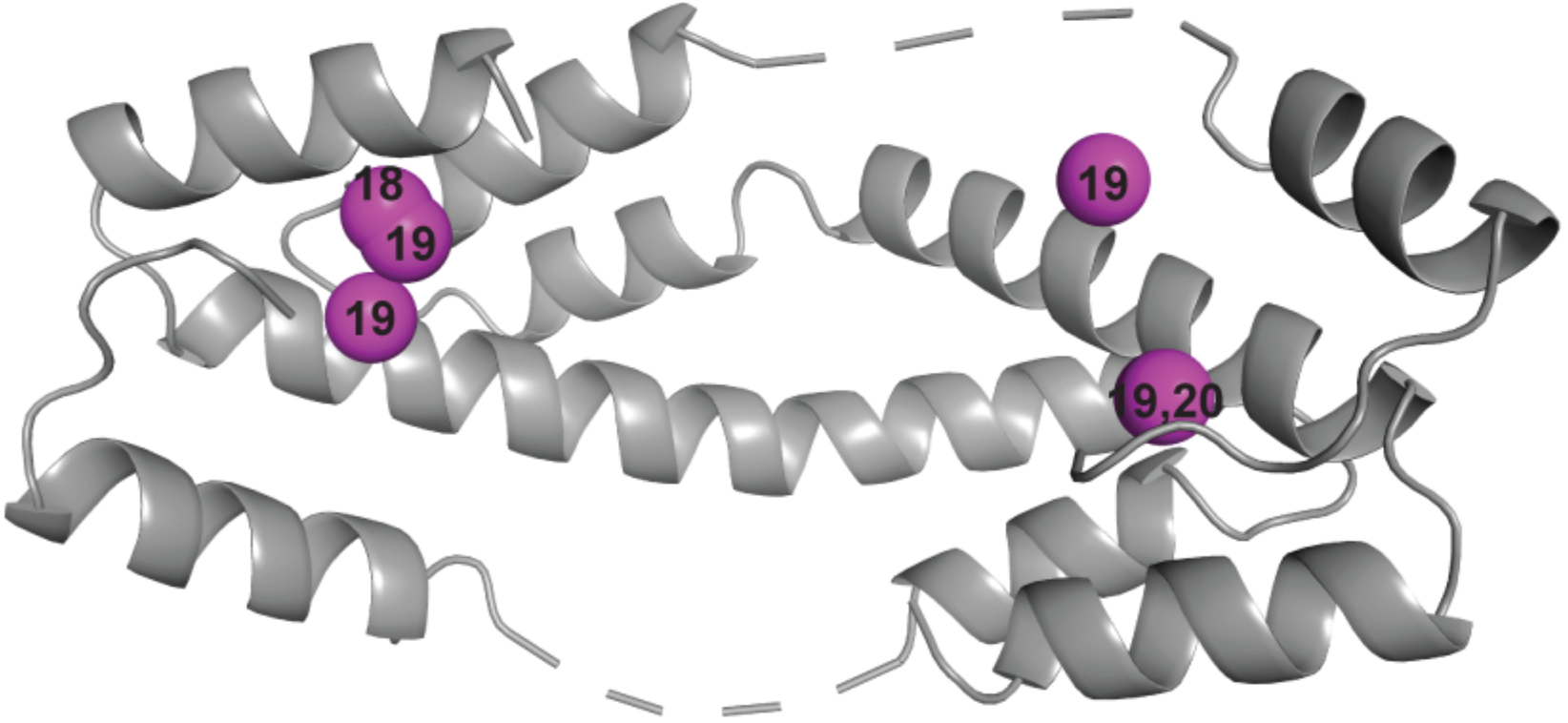
Iodine positions observed in this study, displayed in magenta and labeled by Im7 residue number.

One possible use of anomalous data like those collected here is to perform conformational selections similar to those that are commonly used in NMR and SAXS approaches, and which we recently employed using crystallography data (Salmon *et al.*, 2018; Horowitz *et al.*, 2016). Depending on the quality of the residual electron density from the dynamic components, either an ensemble selection strategy or traditional model building could be aided by anomalous substitutions like those described here.

In our previous publication (Horowitz *et al.*, 2016) on crystals of the Spy-Im7 complex, the anomalous data from the partially occupied Im7 conformations was criticized as being too noisy for analysis (Wang, 2018). In this study, we repeated the measurement of one of the mutants (L19pI-Phe) used in our previous study. Three of the four sites identified here for L19pI-Phe were also observed in our earlier, noisier data (Horowitz *et al.*, 2016). Although our previous cross-validation analyses demonstrated that even our earlier noisier anomalous data contained valuable information (Horowitz *et al.*, 2016; Horowitz, Salmon, *et al.*, 2018) about the Im7 conformations that could then be modeled, the sensitivity of the previously reported experiments was certainly a limiting factor in the technique. In the previous work, the data quality was limited by a combination of detector size and sensitivity due to the CCD detectors employed, as well as air scattering and absorption. The new long wavelength beamline I23 at Diamond Light Source enables the imposition of more stringent criteria for detection of weak anomalous signals. This approach should dramatically improve the ability to delineate dynamic molecules in crystals.

## Acknowledgements

The authors would like to thank C. Travaglini-Allocatelli and A. Di Matteo for useful conversations, Ke Wan for assistance with protein purification.

